# Taguchi Design of Experiment for Optimizing Saline Solution Composition to Increase Electrical Coupling in a Multi-Electrode External EEG Sensor

**DOI:** 10.1101/2025.09.21.677606

**Authors:** Amishi Guliani, Prachi Tyagi, Pawan Tyagi

## Abstract

The effectiveness of noninvasive brain sensing lies in the development of EEG (electroencephalogram) sensors capable of delivering high signal-to-noise ratios for real-time brain-machine interfacing and neuromodulation. A critical factor for achieving this goal is the optimization of electrical coupling between the skin and external electrodes, especially when detecting weak brain signals in the millivolt to microvolt range. In this study, we employed a Taguchi Design of Experiment (DoE) methodology to optimize the composition of an electrolyte solution used to enhance signal quality in a 14-electrode wearable EEG sensor system. We investigated four electrolyte components Sodium Chloride (NaCl), Potassium Chloride (KCl), Sodium Bicarbonate (NaHCO_3_), and Tween-20 surfactant each varied across three concentration levels using an L9 orthogonal array. For each experimental condition, ten measurements were collected to ensure statistical reliability. Two statistical analytical approaches were applied: average signal strength analysis, which exhibited considerable variability and limited diagnostic value, and signal-to-noise ratio (SNR) analysis, which used logarithmic scaling to provide more meaningful insights by minimizing noise effects. Results underscore that while the three salts made comparable contributions, the Tween-20 surfactant was the dominant factor affecting EEG signal strength. Additionally, all four factors exhibited non-linear behavior across concentration levels. This work presents a systematic optimization approach for improving electrolyte formulations and offers practical guidance for enhancing electrical coupling in EEG systems. The results support the futuristic development of high-SNR nanoscale EEG technologies where skin-sensor coupling will be highly sensitive towards saline solution as a conductive medium.

## Introduction

Electroencephalogram (EEG) sensors vary in design and functionality, but a critical distinction lies in whether they require a conductive medium like gel or saline solution to function effectively[1, 2]. Wet and semi-dry electrodes, commonly used in research and medical-grade EEG systems, rely on gel or saline to reduce the electrical impedance between the scalp and the sensor, allowing for more accurate and stable signal acquisition. This is essential for capturing subtle brainwave activity, particularly in high-resolution applications. In contrast, dry electrodes[3], like those found in many consumer EEG headsets (e.g., Emotiv Insight or NeuroSky), sacrifice some signal quality for ease of use and portability[4], as they do not require gel. Semi-dry systems, such as Emotiv EPOC and EPOC X, offer a balance by using saline-based sensors that enhance conductivity while minimizing setup complexity and discomfort. While dry sensors are sufficient for general wellness and basic BCI tasks, gel or saline-based systems remain crucial when precision and reliability are paramount, such as in cognitive research or clinical diagnostics[1].

One of the significant challenges in optimizing EEG signal acquisition with saline solution-based electrodes lies in determining the ideal composition of the saline solution used to improve electrode-skin conductivity [5]. While standard saline typically contains NaCl, researchers have explored the inclusion of additional components [6] such as potassium chloride (KCl), sodium bicarbonate(NaHCO_3_), and surfactants like Tween-20 to enhance ion mobility, maintain pH balance, and improve skin permeability. However, the interplay between these components can significantly affect both the signal quality and long-term stability of EEG recordings. For instance, varying concentrations of NaCl and KCl influence the ionic strength and conductivity of the solution, but can also impact skin irritation and electrode corrosion over time. Sodium bicarbonate helps buffer the solution, maintaining a stable pH conducive to skin health and electrical performance, while Tween-20, a non-ionic surfactant, can reduce skin-electrode impedance by improving wetting and spreading of the solution across the scalp. The challenge lies in balancing these components to maximize EEG signal clarity and understanding the role of each component. Thus, a systematic study of these additives is essential to develop saline solutions that optimize both brain sensor effectiveness and wearability in real-world applications. To address this gap, we systematically varied saline composition using a Taguchi L9 design of experiment[7-9], prepared nine formulations, and evaluated signal quality with a consumer-grade Emotive EEG headset, establishing an efficient framework to identify robust solution chemistries for reliable external EEG recordings.

## Experimental Methods

Four formulation factors were investigated, each at three concentration levels: sodium chloride (NaCl), potassium chloride (KCl), sodium bicarbonate (NaHCO_3_), and a surfactant (Tween 20) [10-12]. Concentrations were expressed in grams per liter (g/L) based on a nominal batch volume of 1 liter. Table 1 provides details of different quantities corresponding to each level of a factor included in this study. The range of levels was determined based on the approximate compositions mentioned. To efficiently assess the main effects and potential interactions among these variables, a Taguchi L9 orthogonal array design was employed. This design enabled the systematic evaluation of four factors at three levels each, using only nine experimental runs. The Taguchi method was selected for its efficiency in reducing the number of experiments while maintaining the ability to identify influential factors and optimize formulation parameters (Table 1).

**Table 1.**
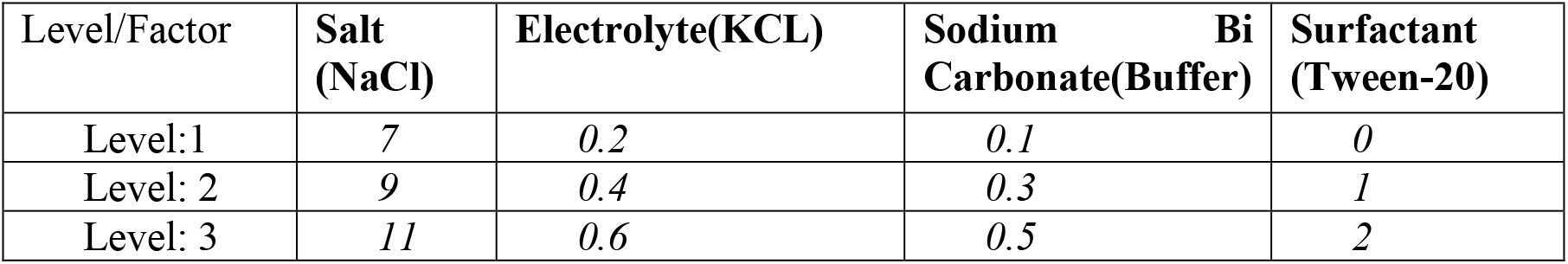
Factors and levels used for the Taguchi Design experiment for saline solution optimization.

| Level/Factor | Salt<br>(NaCl) | Electrolyte(KCL) | Sodium Bi<br>Carbonate(Buffer) | Surfactant<br>(Tween-20) |
| --- | --- | --- | --- | --- |
| Level:1 | 7 | 0.2 | 0.1 | 0 |
| Level: 2 | 9 | 0.4 | 0.3 | 1 |
| Level: 3 | 11 | 0.6 | 0.5 | 2 |

**Table 2.** lists the nine experiments designed based on the Taguchi experimental design approach.

| Table 2: L9 Taguchi Experiment Design for making nine solutions, each with 250 ml volume DI water. |  |  |  |  |
| --- | --- | --- | --- | --- |
| Taguchi Ex# | NaCl (g) | KCl (g) | NaHCO<sub>3</sub> (g) | Tween-20 (g) |
| 1 | 7 | 0.2 | 0.1 | 0 |
| 2 | 7 | 0.4 | 0.3 | 1 |
| 3 | 7 | 0.6 | 0.5 | 2 |
| 4 | 9 | 0.2 | 0.3 | 2 |
| 5 | 9 | 0.4 | 0.5 | 0 |
| 6 | 9 | 0.6 | 0.1 | 1 |
| 7 | 11 | 0.2 | 0.5 | 1 |
| 8 | 11 | 0.4 | 0.1 | 2 |
| 9 | 11 | 0.6 | 0.3 | 0 |

For each experimental run, dry ingredients were added sequentially in the order of the formulation factors— sodium chloride (NaCl), potassium chloride (KCl), sodium bicarbonate (NaHCO_3_), and the surfactant (Tween 20)—into a clean container, followed by the addition of distilled water. To conserve materials, each formulation was prepared at a reduced volume of 250 mL instead of the nominal 1-liter batch. The solutions were gently mixed until all components were fully dissolved. Prior to data collection, freshly prepared EEG electrodes were soaked in the assigned solution for approximately one hour. After preparation, formulations were stored in plastic cups sealed with plastic wrap and kept at room temperature for up to five days.

Recordings were obtained with an EMOTIV EPOC X 14-channel wireless EEG headset with saline sensors[13]. Placement followed the manufacturer’s guidance, with the sensing array positioned on the superior scalp (“top of the head”). Before each trial, sensors were adjusted to part hair away from electrode sites to improve skin contact. Device settings (including placement and acquisition configuration) were held constant across all runs.

All measurements were collected from a single adult male participant to eliminate inter-subject variability. Recording sessions were conducted in the afternoon, with the participant seated and instructed to breathe deeply, keep his eyes open, and minimize both movement and blinking during data collection. For each formulation, three 1-minute trials were recorded. Between trials and across different formulations, electrode positioning and contact quality were checked and readjusted as needed to ensure consistency. Two response metrics were analyzed for each experimental run: (i) the average signal quality metric, aggregated across all EEG channels and trials, and (ii) a larger-is-better signal-to-noise (SNR), used to emphasize robustness. Taguchi analysis was applied to both the response means and S/N ratios to calculate level means, generate main-effects plots, rank formulation factors by maximum–minimum (max–min) delta values, and identify predicted optimal conditions. A one-way ANOVA on the response means was used to estimate the percentage contribution of each factor to the observed variability. Additionally, interactions between factor pairs were evaluated using a severity index (SI%), providing a summary of interaction effects.

## Results & Discussion

After completing the nine experimental runs, the key datasets were compiled and tabulated for further analysis. Table 3 summarizes the results of our experiments from 10 measurements for each experiment. Both average response values and signal-to-noise ratio (SNR) values were evaluated to perform the two types of Taguchi Analysis for an in-depth understanding. We used the bigger the better criterion to perform Average Data and SNR-based analysis (Table 3). These analyses provided insight into the relative influence of each formulation factor and its levels on the measured EEG sensor signal quality.

**Table 3.** 10 Measurement results of 9 different experimental runs and average and SNR for each Taguchi Design of experiment.

| Run# | 1 | 2 | 3 | 4 | 5 | 6 | 7 | 8 | 9 | 10 | Average | SNR |
| --- | --- | --- | --- | --- | --- | --- | --- | --- | --- | --- | --- | --- |
| 1 | 83 | 100 | 92 | 100 | 100 | 67 | 67 | 75 | 92 | 100 | 87.6 | 38.52 |
| 2 | 0 | 0 | 0 | 0 | 0 | 0 | 0 | 0 | 0 | 0 | 0 | -112.78 |
| 3 | 75 | 92 | 75 | 92 | 83 | 92 | 100 | 92 | 100 | 92 | 89.3 | 38.89 |
| 4 | 67 | 58 | 58 | 67 | 58 | 50 | 42 | 50 | 8 | 42 | 50 | 27.24 |
| 5 | 33 | 42 | 25 | 58 | 58 | 25 | 42 | 33 | 8 | 17 | 34.1 | 25.88 |
| 6 | 8 | 17 | 24 | 17 | 8 | 0 | 17 | 17 | 17 | 17 | 14.2 | -94.44 |
| 7 | 33 | 42 | 58 | 33 | 17 | 25 | 42 | 33 | 42 | 58 | 38.3 | 29.95 |
| 8 | 100 | 83 | 58 | 58 | 42 | 42 | 100 | 75 | 75 | 33 | 66.6 | 34.75 |
| 9 | 17 | 25 | 50 | 8 | 8 | 25 | 25 | 17 | 25 | 17 | 21.7 | 23.15 |
| **Average** |  |  |  |  |  |  |  |  |  |  | 44.64 | 1.24 |

Here, we first discuss average-based data analysis using the Taguchi approach. In the first analysis, we focused on understanding the effect of the individual level of each factor by comparing it with the average value (Fig. 1). The first level of NaCl produced an effect greater than the grand average and, therefore, appears to enhance the signal quality compared with the other two levels. Similar to NaCl, the first level of KCl also produced an effect more than the average value. However, for the NaHCO_3_ level 1 and level 3 produced results above the average level and hence improved the signal quality as compared to level 2 (Fig. 3). Interestingly, level 3 of the Tween 20 surfactant produced the highest improvement in the signal quality among all the levels of all the factors; however, level 2 of Tween 20 brought signal quality to the lowest level in comparison with all the levels of all the factors. This study suggests that surfactants are a critical factor. Level 2 of tween 20 being lower than the average data bar suggests that solution coupling efficiency is non-linearly dependent on the surfactant amount in the solution.

**Figure 1.**
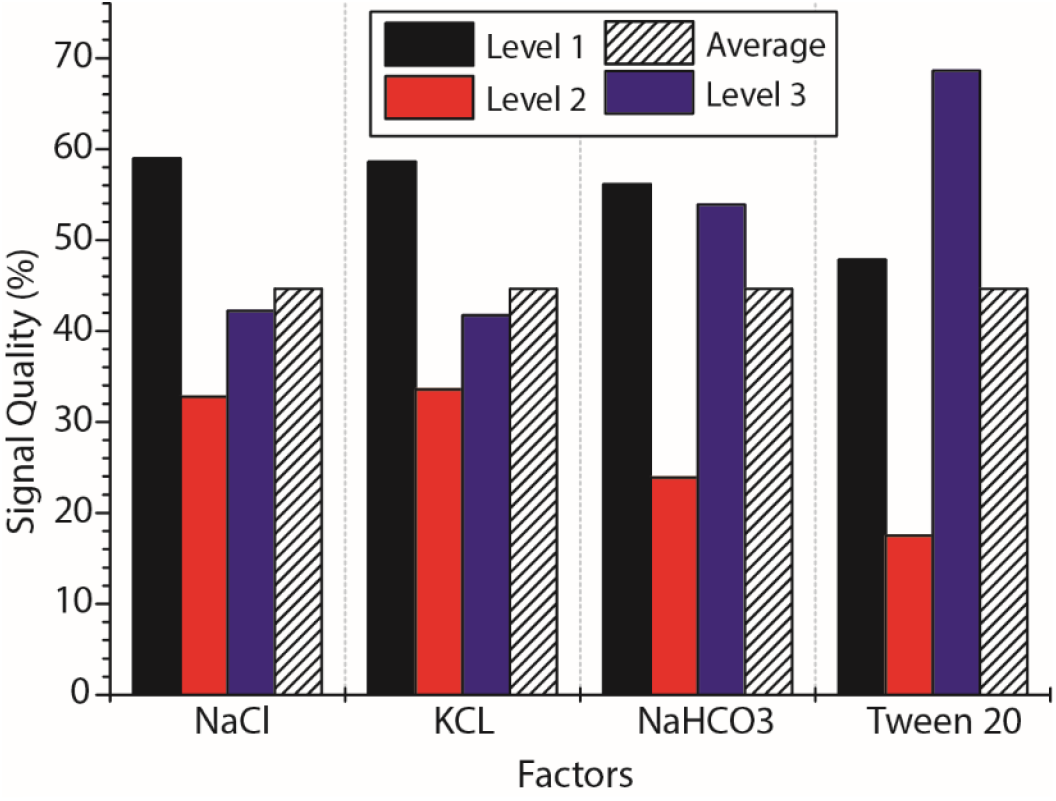
Average analysis of the effect of each level of the four factors.

We also examined whether there was any interaction among the factors used in this study. The Taguchi method provides interaction details using the severity index (%) as shown in Figure 2. NaCl and NaHCO_3_ exhibited the strongest interaction with 69.09%. It means changing the NaCl factor will also require a corresponding adjustment in NaHCO_3_. The interaction between NaCl and KCl amounts interacted with each other with 40.14% SI. The interaction of Tween 20 was significantly higher (37.9%) than observed with NaHCO_3_ (28.94%) and KCl (8.5%). It suggests that Tween 20 surfactants impart different impacts on three salts used in this study. Since Tween 20 and KCl interaction is the lowest, we conclude that these two factors are rather uninfluenced by each other.

**Figure 2.**
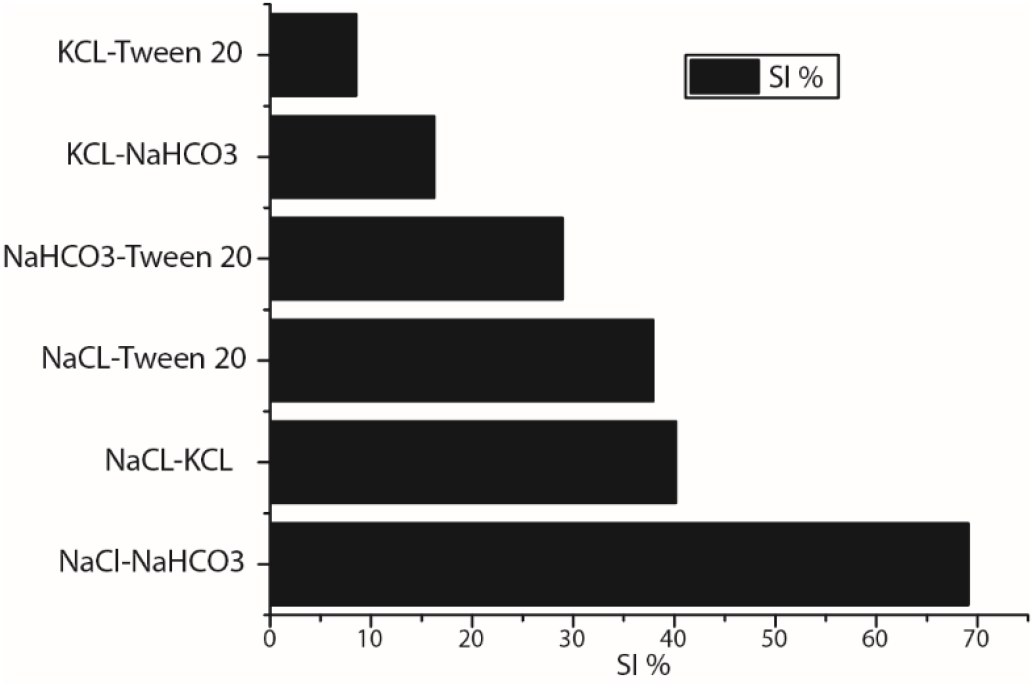
Average analysis of the interactions between factors as severity index (SI) in percent.

**Figure 3.**
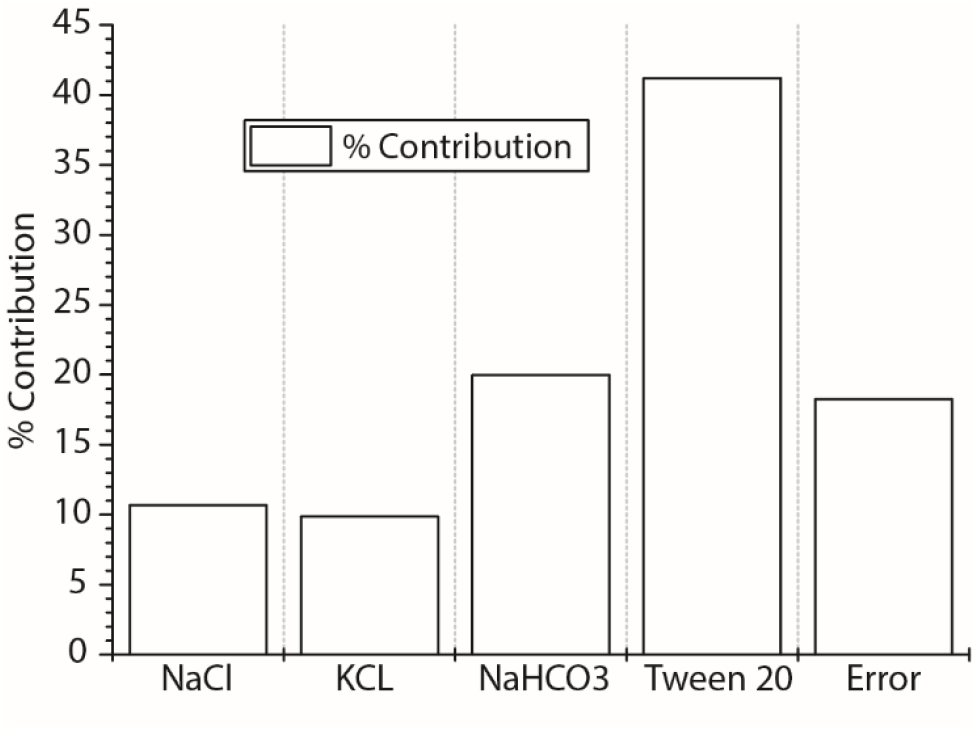
Average analysis of the effect of each of the four factors.

We also performed ANOVA analysis to identify the effect of each factor on EEG sensor signal quality (Fig.3). This bar chart shows how much each ingredient contributed to changes in the average EEG signal quality on a percentage scale. The surfactant (Tween-20) influence on the signal quality dominates at 42%. Sodium bicarbonate is next (20%), while NaCl (11%) and KCl (10%) had smaller effects within the tested ranges. The error term (∼18%) indicates variability not explained by single-factor effects. Since the Contribution of three salts under this average analysis is close to the error contribution, we are unable to ascertain the actual influence. ANOVA analysis also underscores the need to pursue alternative S/N ratio analysis that is better equipped with a mechanism to minimize the error by using the logarithmic scale as compared to the linear scale used in the average analysis.

This main-effects plot shows how each factor level performs relative to the overall average (hatched bar at ∼0): bars above zero indicate better performance for the chosen response, and bars below zero indicate worse performance. For NaCl, Level 3 is clearly above average while Levels 1–2 are below, suggesting higher NaCl within the tested range improved the EEG signal quality response. For KCl, Level 1 is the only positive, indicating that the lowest KCl performed best. Sodium bicarbonate also favors Level 3, with Level 2 markedly detrimental. It is interesting to observe that NaCl salt behaved opposite that of KCl but like NaHCO_3_. For the surfactant (Tween-20), Level 3 (and to a lesser extent Level 1) is positive, whereas Level 2 is strongly negative.

Figure 5 ranks pairwise interactions for the SNR analysis using the Severity Index (SI%). Larger bars mean the two factors do not act additively. The strongest interaction is NaCl × NaHCO_3_ at about 90%, so the robustness of the EEG signal strength depends heavily on how NaCl and bicarbonate are tuned together. Mid-strength interactions are NaCl × KCl (∼49%), KCl × NaHCO_3_ (∼45%), and KCl × Tween-20 (∼43%), which show that the electrolytes influence each other and interact with the surfactant to a moderate degree. Interactions with the surfactant are otherwise weak: NaCl × Tween-20 (∼10%) and NaHCO_3_ × Tween-20 (∼1%). Practical takeaway is that one needs to co-tune NaCl and NaHCO_3_; adjust Tween-20 mostly independently; treat KCl as a secondary knob that interacts with both salts.

**Figure 4.**
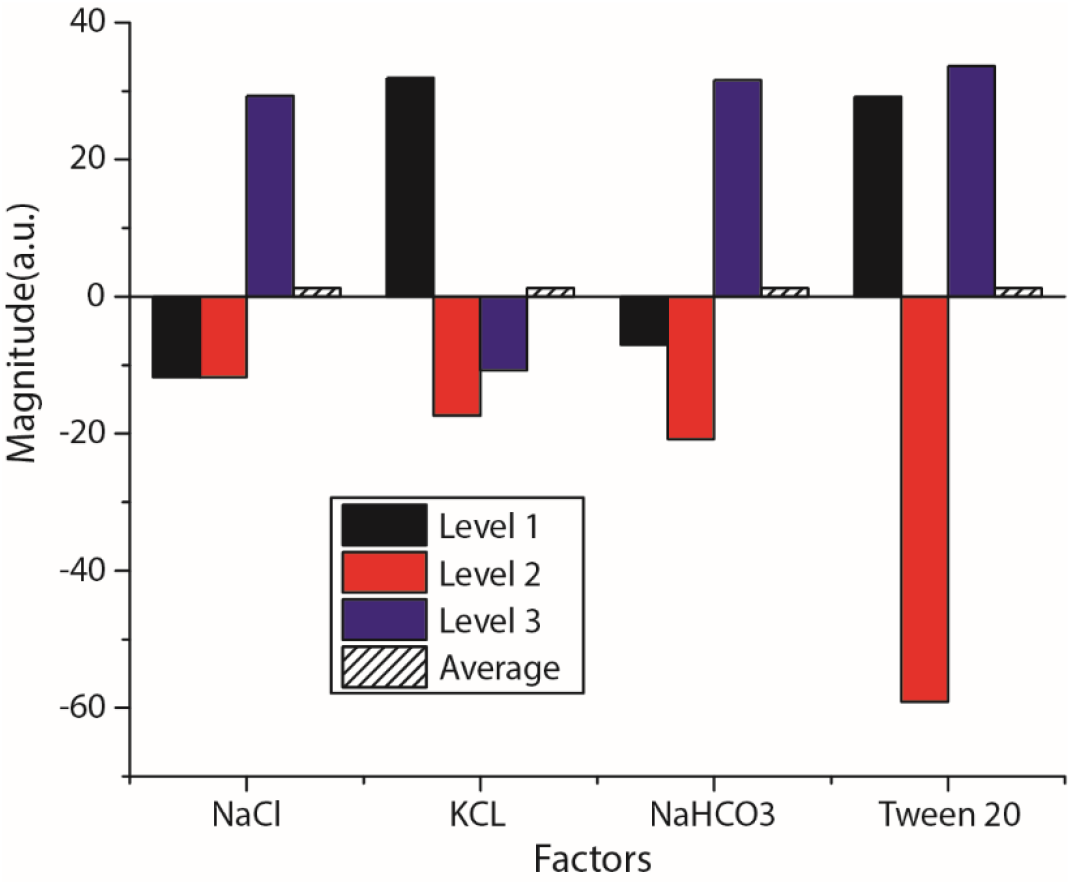
S/N ratio of the effect of each level of the four factors.

**Figure 5.**
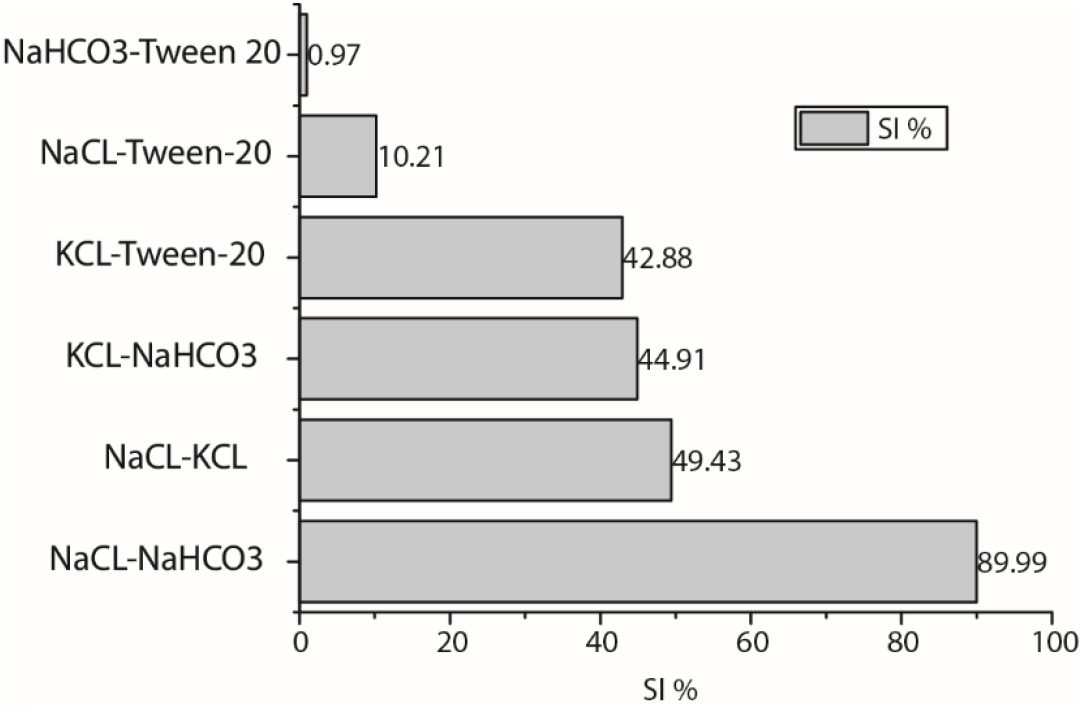
S/N ratio analysis of the interactions between factors as severity index (SI) in percent.

It is interesting to note that the NaCl-NaHCO_3_ interaction was the highest in the average analysis in Figure 2. It is noteworthy that the interaction between pairs of two salts is generally higher than the interaction of Tween-20 surfactant with the salt. The interaction of Tween-20 with KCl is the highest with respect to other salts.

Figure 6 summarizes the ANOVA analysis to identify the impact of individual factors. The bar chart shows each factor’s percent contribution to sensor data. As expected, S/N ratio analysis yielded <2% error. The effect of all the factors is much higher than the error level. Tween-20 contributes the most (about ∼60%), NaHCO_3_ is next (≈ 15%), and NaCl and KCl each contribute less (≈ 10–15%). The Error bar is very small, meaning most of the variability in SNR is explained by these four factors, rather than any additional factor. It is noteworthy that ANOVA analysis under the average analysis method (Fig. 3) is similar to that observed under the SNR analysis approach (Fig. 6). Tween-20 was the most dominant factor in both cases.

**Figure 6.**
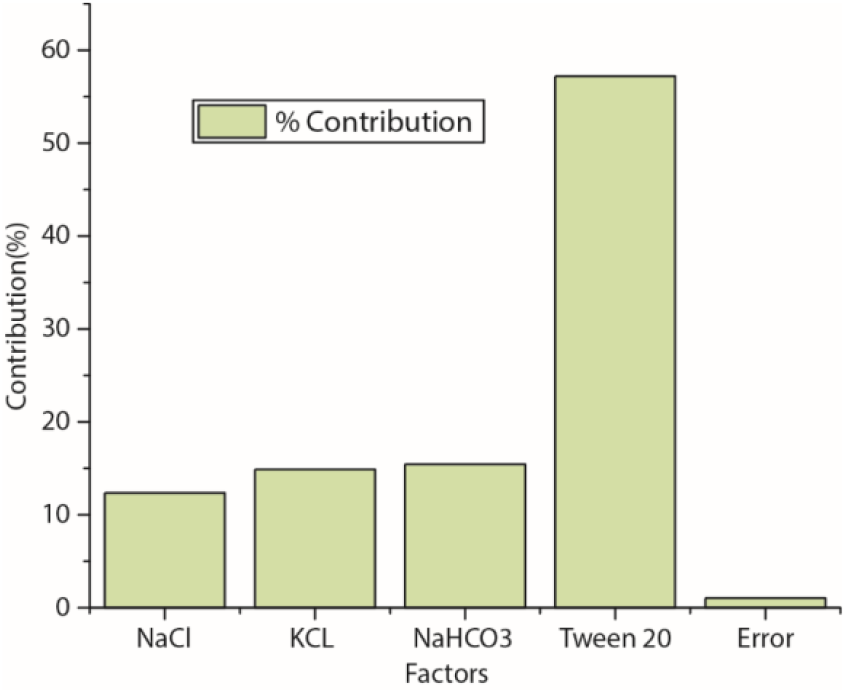
S/N ratio analysis of the effect of each of the four factors.

Table 4 lists the SNR-optimal levels chosen by the Taguchi model: NaCl = Level 3, KCl = Level 1, NaHCO_3_ = Level 3, Tween-20 = Level 3. The “Total Contribution” (121.42) plus the “Grand Average” (1.24) gives the model’s expected result at the optimum (122.66); treating this as a comparative prediction indicates that the optimum combination should yield the most consistent (high SNR performance

**Table 4.** Optimum Results-SNR Analysis.

| Column/Factor | Level | Contribution |
| --- | --- | --- |
| NaCl | 3 | 28.044 |
| KCL | 1 | 30.663 |
| NaHCO3 | 3 | 30.331 |
| Tween 20 | 3 | 32.385 |
| Total Contribution from all factors |  | 121.423 |
| Current Grand Average of Performance |  | 1.24 |
| Expected Result at Optimum Condition |  | 122.66 |

## Conclusions

This study applied a Taguchi L9 design to efficiently identify the dominant formulation factors influencing saline-based external EEG performance. Results demonstrate that surfactant-driven wetting and contact uniformity overwhelmingly determine both mean signal quality and robustness, while bicarbonate serves as a secondary buffer, and salts primarily establish a conductive baseline within the tested ranges. A strong NaCl–NaHCO_3_ interaction highlights the need to co-tune conductivity and buffering, as deviations in one component can compromise performance unless balanced by the other.

The agreement between mean-based and SNR-based analyses strengthens the reliability of the findings, with Tween-20 surfactant emerging as the most critical driver of performance. The L9 framework proved effective in minimizing material use while isolating main effects and revealing interactions. However, limitations include single-participant testing, modest replication, reliance on a proprietary quality metric without impedance, pH, or conductivity validation, and potential carryover effects across formulations. Environmental factors and weighing precision at small scales may also contribute to residual variability. Overall, the results establish surfactant optimization as the key to robust saline formulations for external EEG sensors, with bicarbonate tuning as a complementary lever. Future work should expand subject diversity, employ randomization and counterbalancing, extend recording durations, and incorporate direct impedance, pH, and spectral SNR measurements to strengthen generalizability and mechanistic understanding.

## Author contributions

Amishi Guliani conducted all the experiments, literature review, and wrote the part of the paper. Prachi Tyagi analyzed the results based on average analysis and signal to noise ratio analysis using the Taguchi analysis method and plotted Tables. Pawan Tyagi conceived the concept of the project and advised the experiment.

## Funding

This research was funded by the National Science Foundation CREST Award, grant number HRD-1914751.

### Availability of data and materials

Data included in this paper is available upon reasonable request.

## Declarations

### Conflict of interest

The authors have no competing interests to declare relevant to this article’s content.

## Notes

### Competing Interest Statement

The authors have declared no competing interest.

